# A comprehensive RNA virome from molluscan transcriptomes reveals extensive diversity and modular genome evolution

**DOI:** 10.1101/2025.09.02.673676

**Authors:** Jiaxiang Li, Kewei Chen, Xin Chen, Kaiyang Zheng, Shujuan Sun, Yue Dong, Ziyue Wang, Yixuan Xu, Andrew McMinn, Yeong Yik Sung, Wen Jye Mok, Li Lian Wong, Yantao Liang, Min Wang

**Affiliations:** MoE Key Laboratory of Evolution and Marine Biodiversity, College of Marine Life Sciences, Institute of Evolution and Marine Biodiversity, Frontiers Science Center for Deep Ocean Multispheres and Earth System, and Center for Ocean Carbon Neutrality, Ocean University of China, Qingdao 266003, China; Haide College, Ocean University of China, Qingdao, China; UMT-OUC Joint Centre for Marine Studies, Qingdao 266003, China; Institute for Marine and Antarctic Studies, University of Tasmania, Hobart, TAS, Australia; Institute of Marine Biotechnology, Universiti Malaysia Terengganu, Kuala Terengganu, Malaysia

**Author notes:** Corresponding authors: (YL); (MW).

## Abstract

Mollusca is the second-largest animal phylum and an important marine food resource for humans. While DNA viruses that threaten molluscan aquaculture have received much attention, molluscan RNA viromes remain poorly explored. Here, based on 223 molluscan metatranscriptomes covering eight animal classes, we identified 80 species-level RNA viruses spanning three viral phyla and nine viral families. Phylogenetic result combined structural modeling found a major increase in the number of *Pisuviricota*-related lineages. Extensive modular evolution in viral genomes was observed including gene rearrangements and co-evolution of capsid and RdRP genes. Host prediction linked 78% of the RNA viruses to a diverse range of eukaryotes, including vertebrates, invertebrates, and algae. These findings expand the known diversity of RNA viruses in molluscs and shed light on their phylogenetic relationships, highlighting molluscs as valuable models for RNA virus evolution.

**Graphical abstract:** RNA virus diversity within 223 molluscan transcriptome samples were analyzed based on RdRP identification. Viruses belonging to the phylum *Pisoviricota* were the most abundant; co-evolution between RdRP and capsid proteins of molluscan *Picornavirales* was revealed, and a complex virus-host interaction network was constructed.

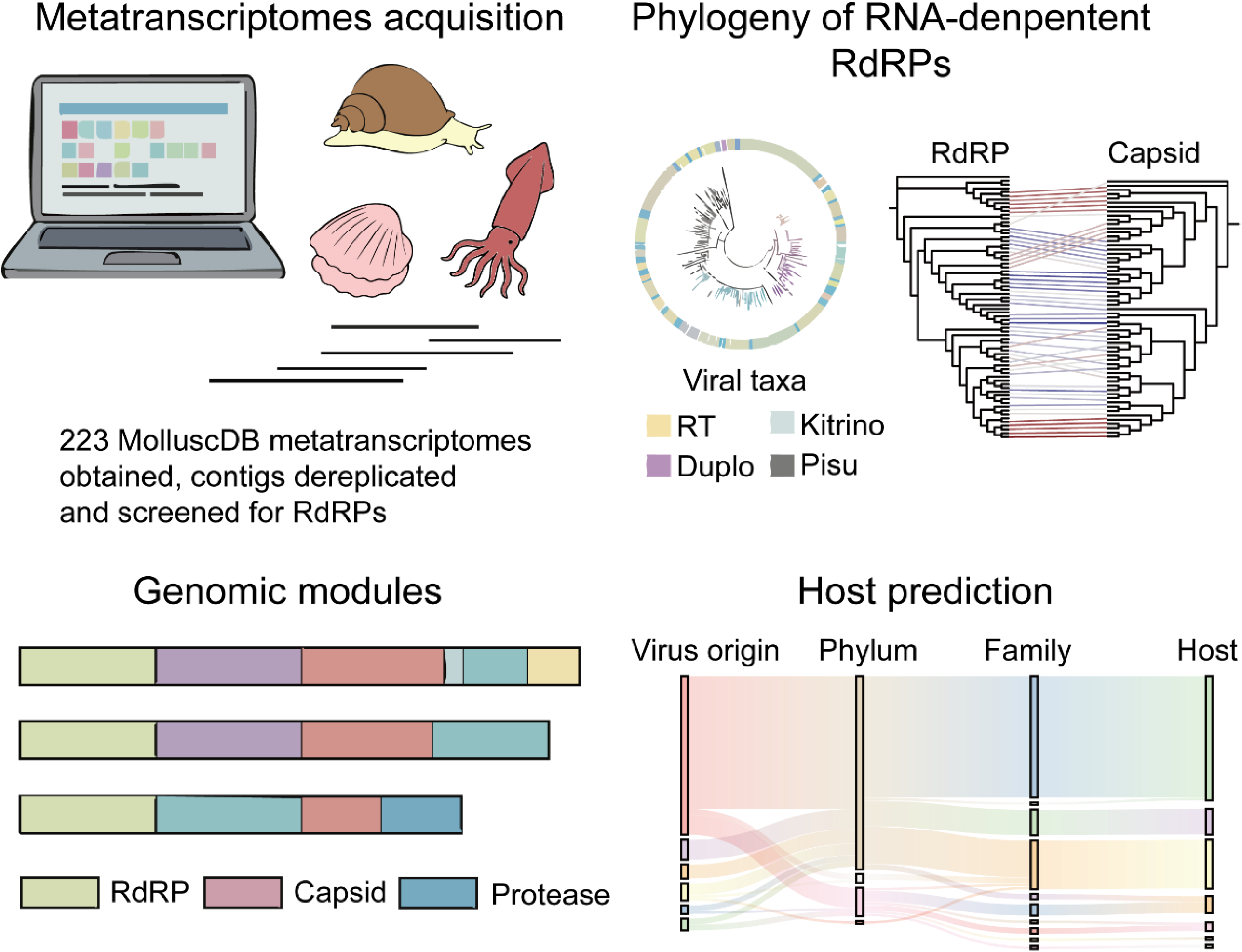

**Highlights:**

1. Comprehensive metatranscriptomic analysis found extensive RNA viral diversity in molluscs.
2. *Pisuviricota* dominate the molluscan RNA virome, substantially broadening known viral lineages.
3. Modular genome evolution involving gene rearrangements and conserved structural features characterize molluscan RNA viruses.
4. Molluscs potentially serve as major ecological reservoirs for diverse RNA viruses infecting various eukaryotic hosts.

## Introduction

Mollusca, which inhabits environments from terrestrial to coastal seas and the deep sea, are the second most diverse animal phylum, with over 100,000 described species [1]. They exhibit high levels of morphological, behavioral, and ecological diversity, which underpin key roles in ecosystems and comprise central positions in food webs. Economically, molluscs account for approximately 27% of global aquaculture production [2]. They also serve as important research models for studies of environmental adaptation [3], nervous system evolution [4], genomic dynamics [5], and disease transmission [6].

Viruses are obligate intracellular parasites and the most abundant biological entities on Earth [7]. Advances in genome sequencing and metagenomics have revealed vast DNA-virus diversity [8], but our understanding of RNA virus biodiversity remains biased toward culturable humans, livestock, and crop pathogens [9]. Metagenomic surveys of invertebrates have found far greater RNA virus diversity than in vertebrates [10]. Shi et al. discovered 1,445 RNA viral genomes by mining transcriptomes from 220 invertebrate species across nine animal phyla. Research on mollusc RNA viruses has lagged behind research on DNA viruses that threaten aquaculture, such as oyster herpesvirus OsHV-1 [11], *Papovaviridae*-like agents [12], and *Iridoviridae* [13], which cause mass mortalities. A small number of RNA virus families (*Togaviridae*, *Reoviridae*, and *Picornaviridae*) have been detected in bivalves [12] through histopathology and electron-microscopy characterization. Recently, 18 and 154 RNA viruses were identified from oysters based on metatranscriptomics, and classified into five families or orders (*Sobelivirales*, *Picornavirales*, *Leviviridae*, *Durnavirales*, and *Yanvirus*) and threee phyla *Lenarviricota*, *Pisuviricota*, and *Kitrinoviricota*, respectively [14, 15]. However, the RNA viruses in other molluscs remains insufficiently explored.

To fill this gap, this study performed a large-scale meta-transcriptomic survey across eight mollusc classes (*Bivalvia*, *Solenogastres*, *Gastropoda*, *Polyplacophora*, *Cephalopoda*, *caudofoveata*, *Scaphopoda*, and *Monoplacophora*) to identify RNA viruses. By mining molluscan viromes and comparing them to public seawater RNA-virus datasets and other known molluscan RNA viruses [10, 14-16], a comprehensive portrait of mollusc-associated RNA viruses containing 80 RNA viral contigs was generated. A systematic phylogeny and host-lineage predictions was also applied to these RNA viruses. Together, the results redefine the molluscan RNA virosphere and shed new light on key patterns and processes driving the evolution of molluscan RNA viruses.

## Methods

### Metatranscriptome acquisition

The identification of RNA viruses was from 223 publicly available, de novo assembled metatranscriptomes obtained from MolluscDB [17]. For detailed information, please refer to Supplemental tables. (If available, please refer to the study where the sample was originally published).

### Primary filtering process

Contigs from MolluscDB were filtered to remove sequences <1,000 nt or encoding rRNAs, then dereplicated at 99 % identity using MMseqs2 easy-linclust [18]. Open reading frames (ORFs) were predicted with Prodigal in “meta” mode [19]. HMMER v3.4 searches [20] against iterative profiles built from known RdRP sequences in NCBI GenBank [21], the Uri Neri et al. Dataset [22], global marine RNA data from Zayed et al. [23], and aquatic virion profiles from Wolf et al [24]. Hits with ≥70 % alignment fraction were realigned with MAFFT v7.475 using default settings [25]. To reduce false positives, we annotated candidates with pfam_scan against Pfam-A (bit-score cutoff 30) and discarded proteins <100 aa or with bit-scores < 30 [26]. Remaining sequences with viral RdRP structural motifs were deemed true positives; lower - scoring hits were manually inspected for conserved RdRP motifs. Finally, PalmScan [27] was used to confirm RdRP domains and to exclude any reverse-transcriptase sequences.

### Contig set augmentation with published genomes

To benchmark the newly predicted viral genomes against known diversity, a “reference set” of published RdRP-encoding sequences was assembled. This collection includes all RdRP hits in NCBI’s NT database plus non - redundant genomes from major RNA virus surveys: the *Leviviricetes* catalog of Callanan et al. [28], the Yangshan assemblage and Wolf et al. [16], *Plastroviruses* described by Lauber et al. [29], the RNA Viruses in Metatranscriptomes (RVMT) database from Neri et al. [22], LucaProt from Shi et al. [30], Oysters RNA data from Wu et al. and Jiang et al. [14, 15] and ocean - derived RdRPs from the Tara Oceans gene atlases [31]. These sequences, together with the novel contigs for downstream phylogenetic and domain analyses, were merged and labeled. All reference and newly generated data are publicly available without restriction.

### Taxonomic annotation of RNA virus

Virus taxonomic assignment used three complementary approaches: (1) the Viral Taxonomic Assignment Pipeline (VITAP) with default settings [32], (2) geNomad with default settings [33], and (3) BLASTp of our RdRP sequences against NCBI’s non-redundant protein database. The results from all three methods were then reconciled to derive consensus taxonomic annotations for the mollusc-associated viruses.

### Phylogenetic reconstruction

The phylogenetic placement of mollusc-associated RNA viruses against the “ reference set” were inferred as follows. First, an environmental RNA virus database under the RCR90 standard (≥90 % pairwise identity, ≥30 % coverage) were screened by BLASTx (e–value 1e– 5) to recruit additional RdRP homologs [34]. Mollusc RdRPs and their reference matches were clustered at 30 % identity via MMseqs2, then subjected to all-vs-all BLASTp (e–value 1e–5) to generate similarity scores. MCL clustering (v.22-282, with an inflation value of 3) [35] was applied to these scores, yielding 33 protein clusters; after removing clusters lacking mollusc RdRPs or containing reverse transcriptases, each cluster was aligned with MUSCLE. For each alignment, HMM profiles were constructed using HH-suite’s hhmake [36], performed HMM–HMM comparisons with hhalign, and converted scores to distances (dAB = – ln[SAB/min(SAA,SBB)]) to produce a 29 × 29 distance matrix. A maximum-linkage dendrogram (R hclust) was cut at height 2.5 to define nine subtrees. Each subtree was realigned with MUSCLE5’s “-super5” option (four guide-tree permutations, three perturbation seeds), then trimmed for ≤20 % gaps with trimAl [37]. Finally, we concatenated all processed RdRP alignments to infer an approximate maximum - likelihood tree in IQ-TREE [38] (1,000 bootstraps), with ModelFinder Plus selecting the best-fit substitution model.

### Three-Dimensional structure network analysis

Representative RdRP structures from each phylogenetic clade were modeled using ESMFold [39]. Pairwise structural alignments were performed with FoldSeek (e–value ≤ 1e–5) [40], including reverse-transcriptase models as outgroups. Alignment bitscores were used to build a 3D-similarity network in Gephi [41] with the ForceAtlas layout. To simplify visualization, the top 100 viruses-associated RdRPs by bitscore plus the outgroups were retained.

### Inference of virus-host interactions

RNA virus hosts were predicted using three complementary methods: (i) CRISPR Spacer Matching. Viral sequences were queried against CRISPR spacers predicted from bacterial and archaeal genomes using BLASTn v2.11.0 (-dust no -word_size 7) [22]. Arrays with ≥3 spacers, each matching a viral sequence with ≤ 1 mismatch and spaced by ≥ 20 bp were retained. Matched repeat sequences were then blasted against IMG/M bacterial and archaeal genomes (perc_identity ≥90, -dust no, -word_size 7) to confirm host candidates and screen for adjacent Reverse transcriptase (RT) or Cas genes. (ii) Taxonomic Inference from Known Virus–Host Pairs. Hosts for annotated RNA viruses were inferred from cross - referencing their VITAP, geNomad, and BLASTn taxonomic assignments against established RNA virus taxa [23]. Putative host lineages were assigned via a Last Common Ancestor approach and validated using ICTV’s curated virus–host records. (iii) Machine - Learning Prediction. RNAVirHost [42], which integrates sequence homology and genome features through a trained classifier, to predict viral host taxa was then applied.

### Comparison of physicochemical property indices of RdRP in RNA viral contigs from different environments

RdRPs from LucaProt, RVMT, and the mollusc dataset were analyzed for five physicochemical properties: hydrophobicity, polarity, relative mutability, transmembrane tendency, and refractivity using the ProtScale server (https://web.expasy.org/protscale/). For each property, scores were computed along the primary sequence with a nine–residue sliding window. Positional fluctuations in each individual index are presented as line graphs, and comparisons of the full set of physicochemical profiles across the three RdRP groups are summarized in violin plots generated in GraphPad Prism 8.

### Estimation of the relative abundance of RNA viruses

To evaluate the prevalence of RNA viruses identified in molluscs, qualified reads were mapped to RdRP-bearing contigs using Bowtie2 (v 2.5.4) [43] with the modified “very sensitive” option (–local -D 20 -R 3 -N 0 -L 16 -i S,1,0.50 -I 0 -X 2000 –non-deterministic) [23]. To minimize potential bias in abundance estimation caused by overrepresented polyA stretches, both the reads and viral sequences were trimmed at polyA or polyT ends using BBDuk (v37.62) (https://jgi.doe.gov/data-and-tools/bbtools/), without iterative trimming, as the second round removed no additional polyA/polyT reads. Mapped reads were converted into sorted BAM files using SAMtools (v1.21) [44], which were then used to calculate overall genomic coverage and abundance. Read mappings were subsequently filtered at thresholds of ≥90% nucleotide identity and ≥75% read length coverage using CoverM (v0.7.0) (https://github.com/wwood/CoverM). Transcripts per million (TPM) were calculated with the “-m tpm” option in CoverM to provide an estimate of relative abundance. While the conventional approach of identifying expressed genes by comparing metagenomic and metatranscriptomic signals is not applicable to RNA viruses due to their RNA-based genomes, the activity of the RNA viruses assessed solely based on metatranscriptomic coverage and depth thresholds. Specifically, RNA viruses with ≥50% of the genome covered at 1× or more with a median coverage depth > 0 on the coding strand were classified as actively transcribing, as defined by the Coclet et al [45] transcriptional activity criteria.

### Statistical analysis

All statistical analyses were carried out in R v4.3.2. Alpha- and beta-diversity metrics, rarefaction curves, and scatter plots were generated with ggplot2 v3.4.4 and ape v5.7.1. Violin plots were drawn in Python using seaborn and matplotlib.pyplot. Protein–structure bitscores were used to build a 3D-similarity network in Gephi via the ForceAtlas layout. Phylogenies were visualized in iTOL [46], and virus–host Sankey diagrams were created with SankeyMATIC (https://sankeymatic.com).

## Results

### RNA viral contigs in molluscs

Samples collected from a wide range of molluscs (Fig. 1A), collected 223 molluscan metatranscriptomes spanning eight classes (*Bivalvia*, *Solenogastres*, *Gastropoda*, *Polyplacophora*, *Cephalopoda*, *Caudofoveata*, *Scaphopoda*, and *Monoplacophora*), with bivalves comprising the largest class (48.9%, Fig. 1A). The assembly of these metatranscriptomes yielded 9,876,760 contigs ( ≥ 1 kb) and 12,086,615 predicted proteins. Using our custom pipeline (Fig. S1), we iteratively searched for RdRP domains and identified 2043 candidate RNA virus contigs from 35 of the 223 metatranscriptomes. Pfam_scan and PalmScan filtering reduced these to 80 high confidence RNA viral contigs, whose RdRP lengths ranged from 350 to 1,400 aa and GC contents from 30 % to 55 % (Figs. 1C).

**Fig 1.**
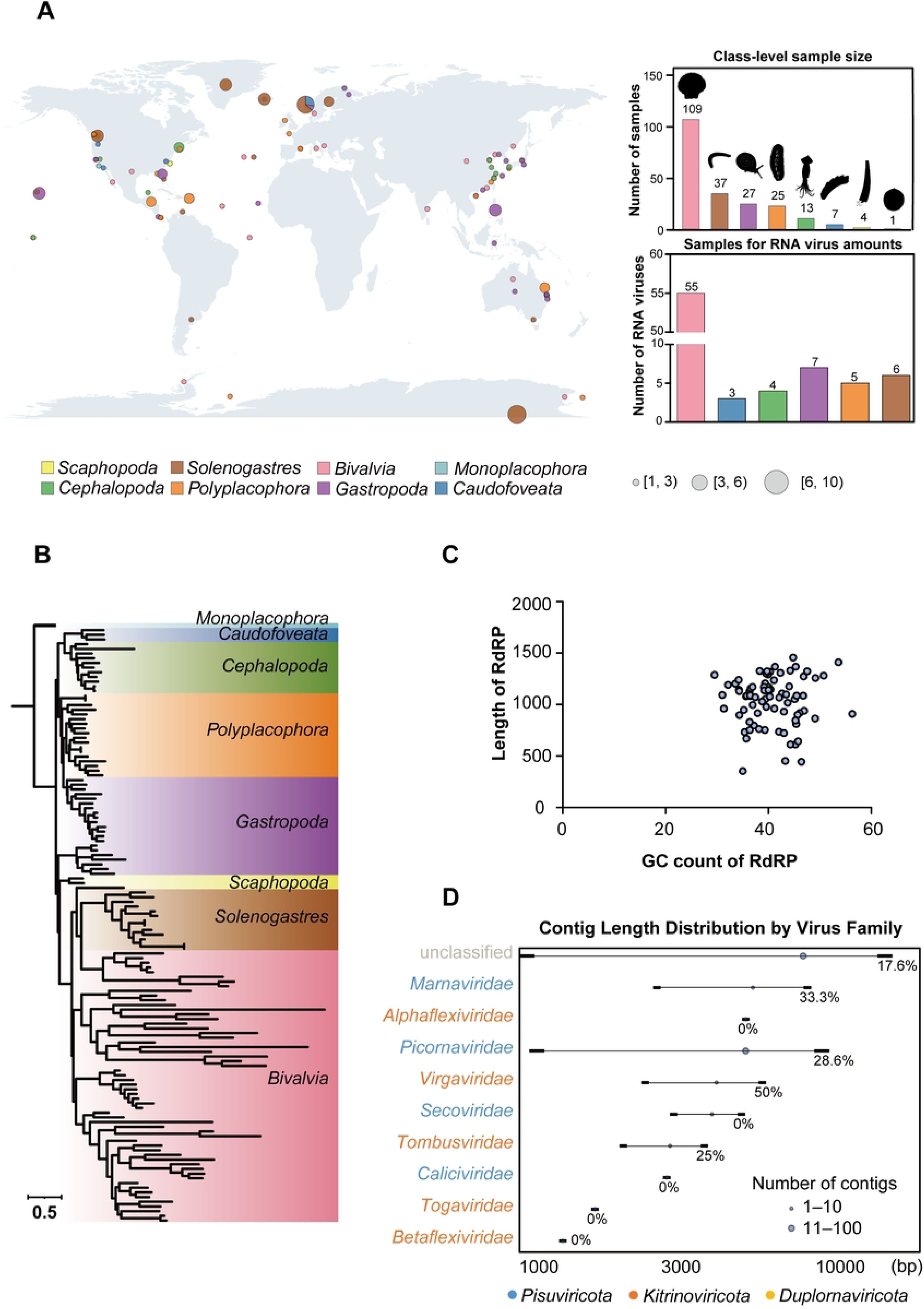
The distribution of molluscan metatranscriptomic data analyzed in this study. (A) Overview of the samples analyzed in this study. Pie charts show the composition of mollusc species collected at each location. The total area of each pie chart is proportional to the number of individuals collected. Colors represent different mollusc species, only samples with clear geographic information are shown. (B) phylogenetic trees of some mollusk samples were constructed using maximum likelihood (ML) based on nucleotide sequences of the COI gene. The color scheme matches Fig 1A. (C) The length of RdRP and GC content identified in this study. (D) Distribution of 80 viral contigs by length, taxonomic family. Segments represent the spread between the minimal and maximal lengths of contigs assigned to the taxonomic family rank. Dot size represents the number of contigs within the taxonomic family rank.

Taxonomic annotation via VITAP, geNomad, and BLAST against the nr database assigned these RNA viruses to three viral phyla, including *Pisuviricota* (n= 66, 82.5 %), *Kitrinoviricota* (n= 10, 12.5 %), and *Duplornaviricota* (n= 1, 1.5 %), positive-sense single-stranded RNA (+ssRNA, n= 76) viruses are the dominant RNA viruses. In addition, three RNA viruses could not be classified (Fig. 1D). At the family level, 21.3 % of RNA viruses (n= 17) remained unclassified, though 12 of these unassigned sequences are likely *Pisuviricota* lineages at phylm level. When the data extracted here was compared with the invertebrate mollusc RNA viruses dataset [10], it was found that both studies recovered members of *Pisuviricota* and *Kitrinoviricota* (Fig. S2). Unlike previous studies, *Negranaviricota* and *Lenarviricota* viruses were not identifies here. In addition, a *Duplornaviricota* virus in *Proneomenia*, classified as *Ghabrivirales* was detected, which has not been reported in previous molluscan virus studies.

### RdRP Phylogeny and expansion of molluscan RNA viral diversity

Molluscan RNA viruses displayed broad phylogenetic diversity when classified using the RdRP marker gene. Rooting the phylogenetic alignmenst on reverse transcriptases, the global RdRP tree splits into four major clades—Reverse Transcriptase, *Kitrinoviricota*, *Pisuviricota*, and *Duplornaviricota* (Fig. S3). While each viral phylum formed its own independent branch, the unclassified sequences were intermixed with *Kitrinoviricota* and *Pisuviricota*, potentially revealing a novel viral clade. To resolve in-depth phylogenetic relationships, an iterative maximum - likelihood workflow based on 2,106 RdRP sequences plus 69 RT sequences as outgroups was performed (see Methods), yielding 5 viral clades associated with the virus found in this study. Subtree analyses (Fig. 2) showed many mollusc-derived viruses form distinct, well-supported clades that diverged from previously recognized supergroups [10]. Notably, the Picorna, Toli, Martelli, Amarillo and Ghabri lineages, once only sparsely sampled now encompass substantially expanded clusters (Fig. 2), underscoring the previously underappreciated phylogenetic breadth of RNA viruses in molluscs [14, 15].

**Fig 2.**
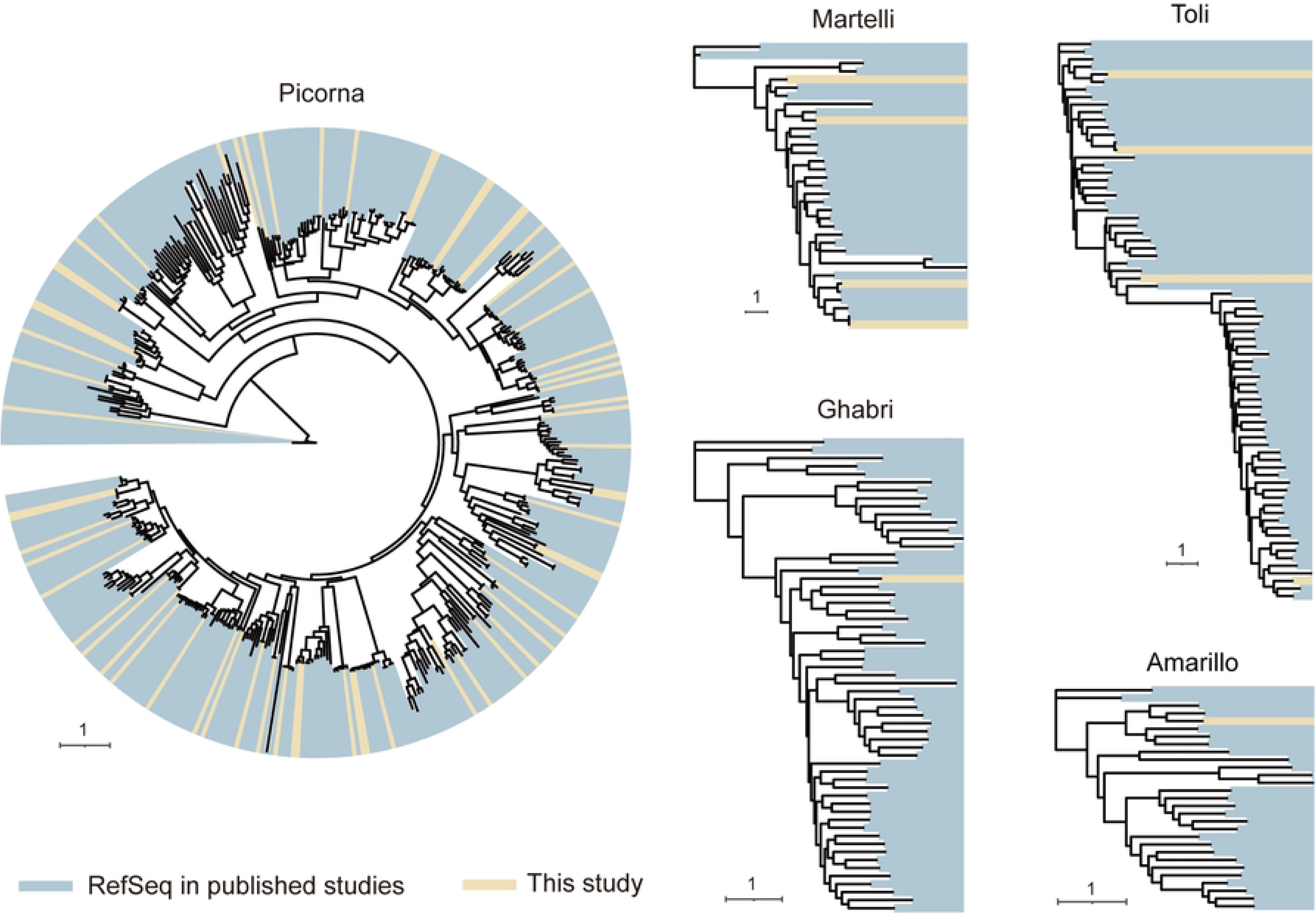
Phylogenetic diversity of five RNA viral supergroups containing molluscan viruses. Each phylogenetic tree was estimated using a maximum likelihood method based on an amino acid alignment of the RdRP domain. Newly identified viruses are marked in yellow, and viral RdRP sequences from published metagenomic studies are marked in blue. The name of each supergroup is shown at the top of each phylogeny.

### Three-dimensional structural clustering network based on RdRPs

The RdRP gene plays a crucial role in the replication and transcription processes in a range of organisms [47, 48]. Amino acid residues involved in key processes, such as catalysis and nucleotide selection, are often highly conserved [22]. In this study, the RdRP sequences derived from the samples, reference RdRP sequences, and RT sequences were subjected to three-dimensional structural clustering. The top 100 sequences with the highest bit scores were selected for visualization. The clustering results based on three-dimensional structures revealed that each viral order could be independently clustered. However, a few viral orders were not entirely distinguishable in their three-dimensional structures, likely because certain sequences within these orders resembled those of other orders (Fig. 3). This observation aligns with findings from the phylogenetic tree analysis (Fig. S3).

**Fig 3.**
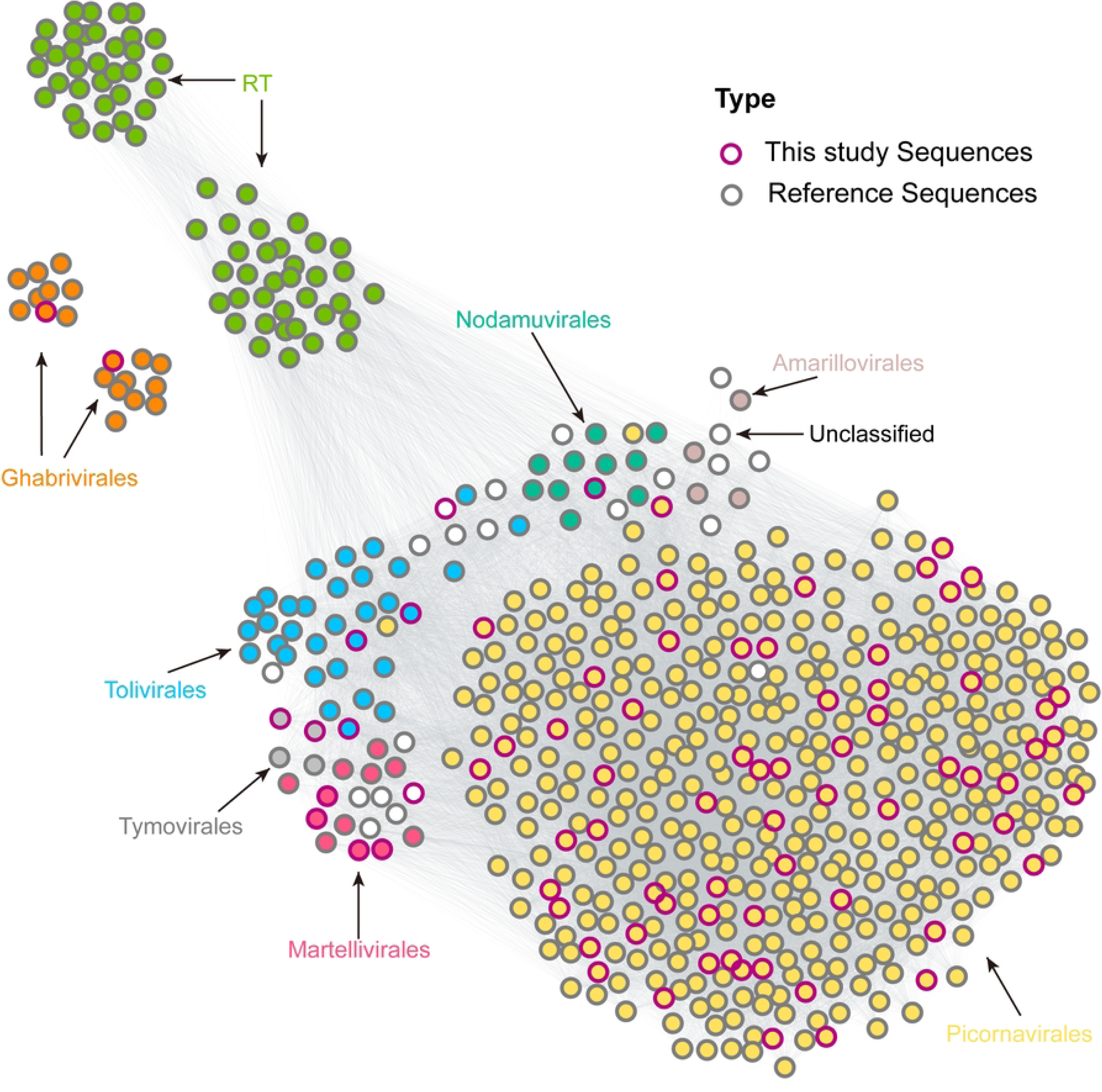
RdRP-based structure network analyses of molluscan RNA viral contigs. Three-dimensional structure similarity network of RdRP and Reverse Transcriptase protein domain structures. Each node represents a different structure. For clearer visualization, the top 100 best hits associated with each node are selected as significant nodes, with structure bit as the weight. Different colored nodes represent the commented virus order.

### Exploration of potential hosts of RNA viral contigs

Viruses significantly influence a wide range of microbial-mediated processes through their interactions with host organisms. To identify the hosts of RNA viruses in the mollusc samples, three approaches were employed: comparison of RNA viral contigs with predicted bacterial and archaeal CRISPR spacer sequences, utilization of available host information for established viruses, and application of the RNAVirHost tool [42]. However, comparative analyses between CRISPR spacers and RNA viruses yielded no significant alignment results. This outcome can primarily be attributed to the inherent limitations of using CRISPR spacers to investigate RNA viruses infecting prokaryotic organisms [49]. RNAVirHost provides a cost-effective and efficient strategy for host prediction. This study combines the above methods to assist RNA virus analysis, aiming to obtain a more accurate viral host lineage. The results indicate that 78% of the RNA viruses were associated with potential hosts. To date, the majority of RNA viruses have been linked to eukaryotic hosts. Most of the potential hosts are eukaryotes, with the majority of RNA viruses infecting *Gnathostomata*, Invertebrates, and Stramenopiles (Fig. 4). The *Picornaviridae* were the most abundant family, with primary hosts in mammals, birds, and insects, supporting broad virus–eukaryote associations of molluscan RNA viruses.

**Fig 4.**
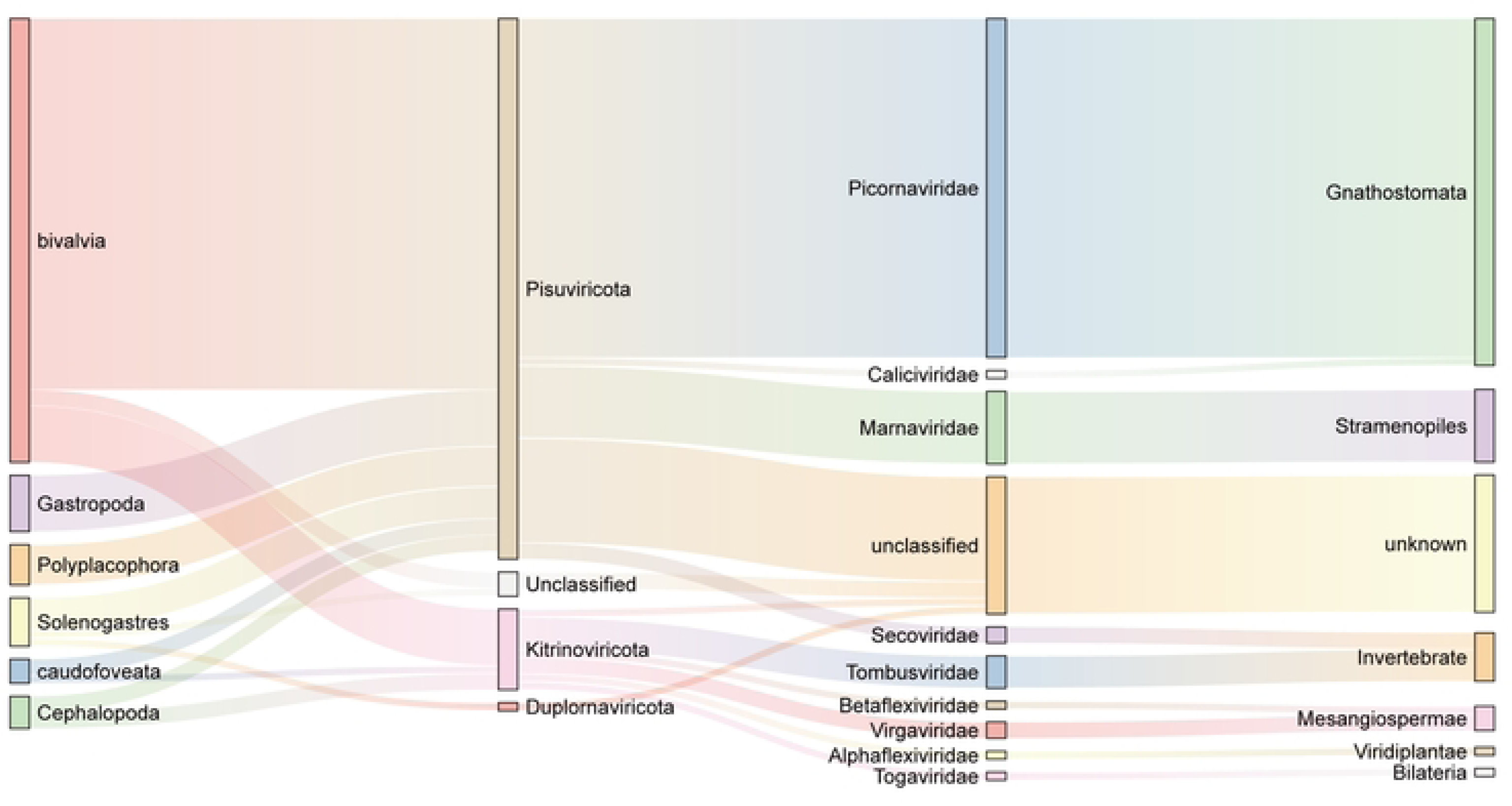
The connectivity between molluscan RNA viral contigs and potential hosts. Sankey diagram showing the relation among the sample group of RNA virus origin, the annotation of RNA viral contigs and the predicted hypothetical host.

### Patterns of RNA virus genome evolution and Protein features

Viral genomes from key clades were examined and extensive modularity, including segment fusions and protein rearrangements was observed (Fig. 5A). RdRP-encoding segments ranged broadly in length but clustered around 1,200 nt. Whereas *Tombusviridae*, *Virgaviridae*, *Picornaviridae*, and *Marnaviridae* typically separate capsid protein (CP) and RdRP on different segments, instances where both genes co - occur on the same segment were identified here. Several examples of non-homologous replacements of structural modules were also found: one unknown-family RNA virus contig carried a CP similar to *Marnaviridae* and Dicistro_VP4, while another swap replaced a CP with an IgA - protease–like domain, eliminating the CP entirely. In *Picornaviridae* RNA viruses, distinct CP variants nonetheless shared high 3D - structural similarity (Fig. S4A) and conserved β-sheet and loop motifs (Fig. S4B), supporting a common fold despite sequence divergence. Single-jelly-roll capsid proteins (SJR CPs) are often found in basal lineages, so these early viruses likely encoded only an RdRP and an SJR CP. Most eukaryotic RNA viruses probably evolved from these simple ancestors [50].

**Fig 5.**
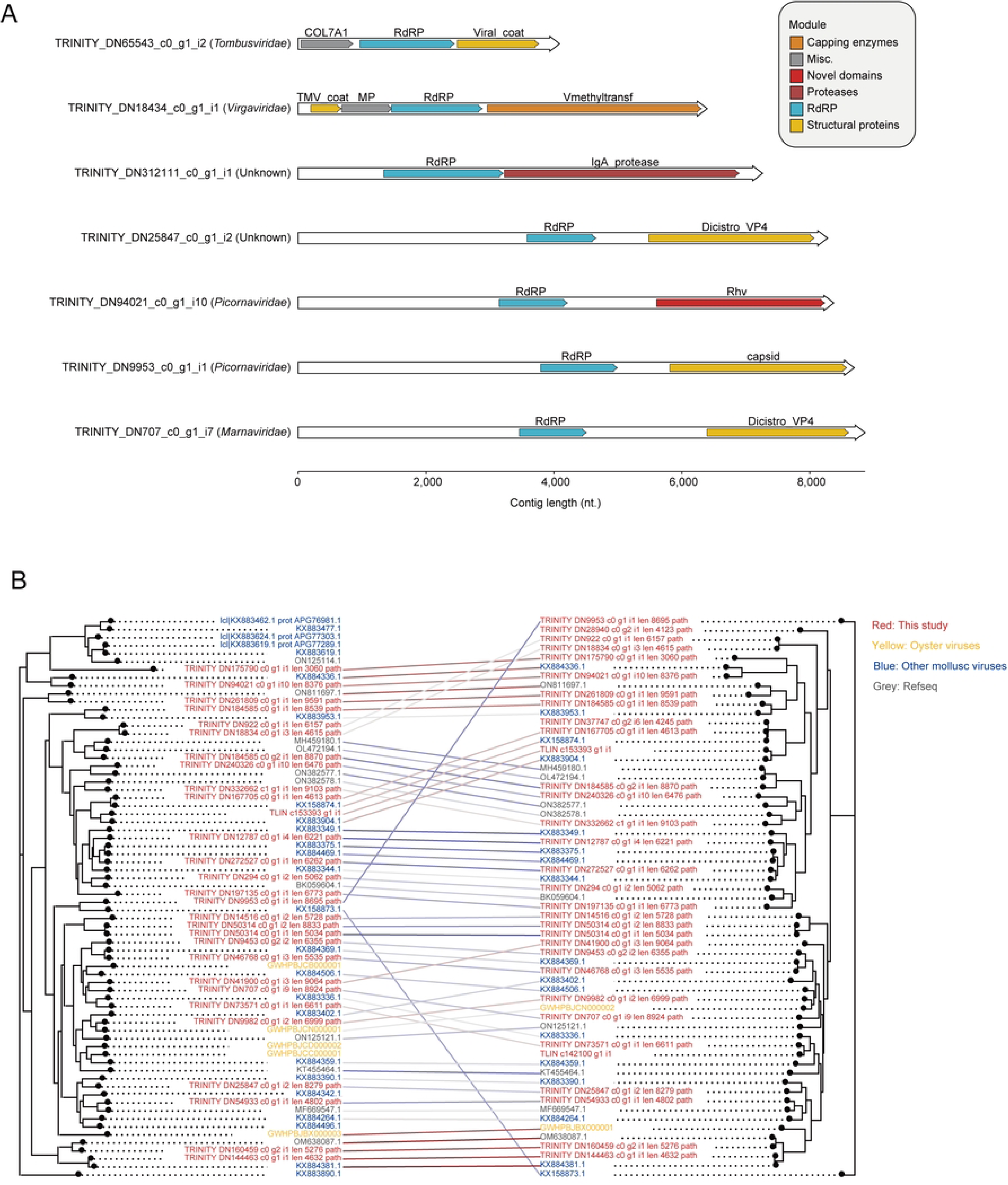
Genomic features and phylogenetic comparison of molluscan RNA viral contigs. (A) Homologous domains are shown as boxes of the same color (see key on the right). Domains not commonly found in RNA viral contigs are shown in red and are labeled above the corresponding boxes. Virus taxa and contig identifiers are noted to the left of each virus genome. At the bottom, scale indicating the length in nucleotides. (B) Comparison of the topologies of RdRp- (left) and capsid protein-based (right) phylogenetic trees of picornaviruses identified in this study. Trees were inferred with IQ-TREE v2.4.0 using ModelFinder (MF) and 3,000 ultrafast bootstrap replicates; bootstrap values > 70 are shown. Tip labels are color-coded by data source.

*Picornavirales* are the most abundant RNA viruses in coastal waters [51] and several mollusc-associated members were identified in this study (Fig. 5B). Most were closely clustered with Beihai picorna-like virus, Wenzhou picorna-like virus and Bivalve RNA virus G5, whereas one picorna-like virus (TRINITY_DN175790_c0_g1_i1_len_3060) was markedly divergent. Comparison of RdRP and capsid-based phylogenies of picornaviruses revealed no evidence of recombination among major clades, yet significant topological incongruence between the RdRP and capsid trees was observed in several branches, suggesting that their capsids may have been lost and subsequently reacquired from other sources.

To characterize RNA virus protein features in molluscs, we analyzed RdRP sequences from the *Pisoviricota* evolutionary branch (Fig. S5). Comparative analysis of physicochemical properties, including hydrophobicity, polarity, mutability, transmembrane propensity, and refractivity revealed distinct patterns between mollusc-derived RNA viruses, LucaProt, and RVMT (Fig. 6). Notably, molluscan RdRPs exhibited statistically significant differences in refractivity and mutability compared to both LucaProt and RVMT (Student’s t-test, p < 0.05). In contrast, no significant divergence was observed in polarity or transmembrane propensity across these groups.

**Fig 6.**
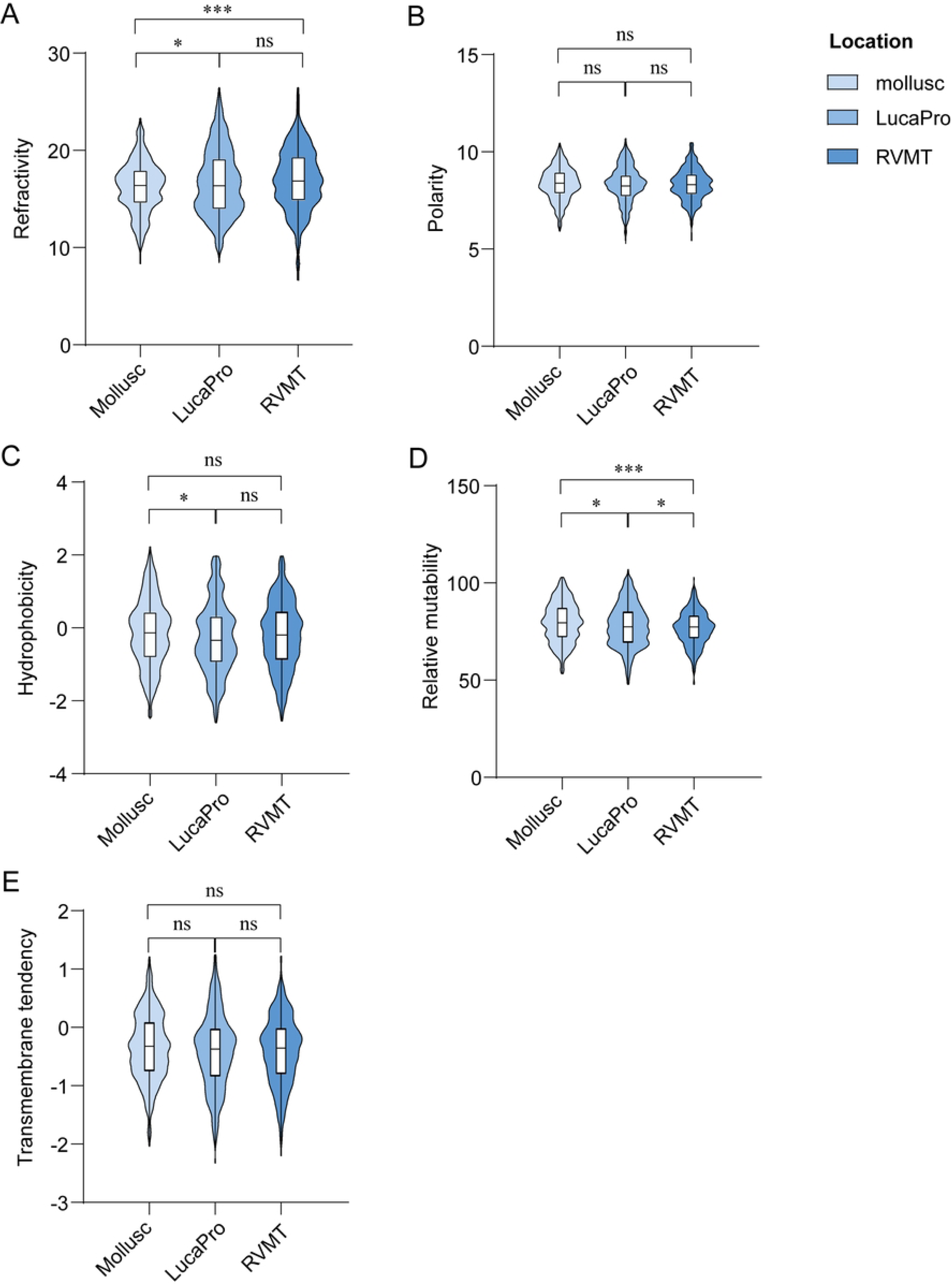
Comparison of physicochemical property indices of proteins from molluscan RNA viral contigs with those from LucaProt and RVMT. (A) The comparisons of refractivity of RdRPs proteins among different habitats. (B) The comparisons of polarity of RdRPs proteins among different habitats. (C) The comparisons of hydrophobicity of RdRPs proteins among different habitats. (D) The comparisons of relative mutability of RdRPs proteins among different habitats. (E) The comparisons of transmembrane tendency of RdRPs proteins among different habitats.

### Relative abundance and activity of RNA viruses

The relative abundance of RNA viruses exhibits considerable variability (Fig. S6A). The log-transformed TPM values indicated that 21 viruses exceeded the threshold of 6. This indicates that the reads corresponding to these viruses comprise at least 10% of the total aligned reads. The 21 viruses belong to four families: *Marnaviridae*, *Virgaviridae*, *Picornaviridae*, and *Alphaflexiviridae*. Among the remaining five viral lineages, plant-infecting families predominated, particularly *Tombusviridae*, *Betaflexiviridae*, and *Secoviridae*. Picorna-like RNA viruses have previously been reported to infect mollusks [52], and a calicivirus (TRINITY_DN175790_c0_g1_i1) was recently identified in the freshwater bivalve *Congeria kusceri* (*C.kusceri*), a family that also includes major human gastrointestinal pathogens [53, 54], providing additional support that *C. kusceri* may serve as a potential calicivirus host. However, this contig accounted for less than 0.6% of total reads, indicating limited environmental distribution or low persistence in *C. kusceri* relative to the highest-abundance viruses.

Metatranscriptomes offer a unique opportunity to further understand which viruses are active, strand-specific mapping information was leveraged to identify actively replicating viruses on the basis of detecting both coding and non-coding genome strands. This enabled a comparative analysis of activity levels for RNA viruses in mollusc. RNA-virus activity was detected across the surveyed molluscs (Fig. S6B). *Picornaviridae* were active in *Congeria kusceri*, *Tonicella lineata*, *Pholidoskepia*, *Proneomenia sp.*, *Yoldia eightsii*, and *Chrysomallon squamiferum*. *Marnaviridae* were active in *Congeria kusceri*, *Tonicella lineata*, and *Pholidoskepia* (Supplemental tables). In strand-specific datasets, concurrent detection of coding and non-coding strands further supported active replication. Collectively, picornaviruses appear widespread and frequently active across molluscs, while marnaviruses detected in benthic grazers most likely reflect algal infections present in diet or biofilms.

## Discussion

This study examines the diversity and evolutionary features of RNA viruses through an in-depth analyzsis of 232 molluscan metatranscriptomes, spanning eight classes, significantly expanding the known RNA viral diversity of molluscs [55]. Phylogenetic reconstruction showed that most new viruses formed distinct clades, divergent from established taxa (Fig. 2) [10], suggesting that molluscs may serve as critical reservoirs for RNA virus evolution. Three-dimensional structural clustering of RdRPs corroborated phylogenetic relationships, demonstrating conservation at the viral order level (Fig. 3), though structural overlaps in minor clades may reflect evolutionary convergence or horizontal gene transfer.

Comparison with earlier studies underscores the novelty of our dataset (Fig. S2), while both our work and previous invertebrate surveys recovered *Pisuviricota* and *Kitrinoviricota*, our larger-scale analysis across 223 metatranscriptomes identified 80 high-confidence RNA viruses, providing a standardized baseline for cross-library comparisons. Notably, we detected a *Duplornaviricota* (*Ghabrivirales*) lineage from *Proneomenia* that has not been documented in molluscs, and observed a strongly skewed composition dominated by *Pisuviricota* (∼82.5%). These additions extend both the breadth and depth of molluscan RNA viromes and provide a new phylum-level dimension that was missing from previous records.

Modular evolution of RNA viral genomes was prominent, particularly in the organization of capsid protein and RdRP coding regions [22]. Instances of CP and RdRP co-encoded on a single genomic segment were observed in families such as *Tombusviridae* (Fig. 5A), alongside non-homologous structural gene displacement, supporting the hypothesis of modular recombination as a driver of host adaptation [22]. For example, while CP sequences in *Picornaviridae* exhibited variability, their three-dimensional structures remained highly conserved (Fig. S4A), indicating functional constraints despite sequence plasticity. These findings align with the SJR CP origin hypothesis [50], that proposes that eukaryotic RNA viruses likely arose from ancestors encoding RdRP and simplified CPs.

From an ecological perspective, host prediction linked 78% of RNA viruses to eukaryotic organisms (e.g., plants, algae, vertebrates, invertebrates, Fig. 4), potentially due to RNA viruses primarily targeting eukaryotic hosts [56]. RNA viruses that infect protists have been found in *Congeria kusceri*, *Meretrix petechialis*, *Panopea generosa*, *Perna viridis*, *Pinctada maxima*, *Reishia clavigera*, and *Tonicella lineata*. Plant-infecting RNA viruses have been detected in *Mimachlamys nobilis*, *Tridacna squamosa*, and *Euprymna berryi*. This suggests that molluscs may act as ecological bridges for viral transmission by filtering and accumulating viruses from the environment. In future, integrating cross-host metagenomics spanning diverse molluscs, *in vitro* assays, and single-cell technologies will verify mollusc-virus interactions and their ecological and evolutionary implications.

## Acknowledgments

We appreciate the computing resources provided byCenter for High Performance Computing and System Simulation, Laoshan Laboratory, the High-Performance Biological Supercomputing Center at the Ocean University of China, the Marine Big Data Center of the Institute for Advanced Ocean Study of the Ocean University of China, and the IEMB-1, a high-performance computing cluster operated by the Institute of Evolution and Marine Biodiversity.

## Author contributions

YL, AM and MW: conceptualization, revision, project administration, supervision, and funding acquisition. JL: methodology, visualization, writing, and original draft preparation. KC: writing-review, investigation. XC and KZ: software, validation, and visualization. SS: formal analysis, interpreted data. YD and ZW: methodology, script. YYS, LLW and WGM: revision. The authors read and approved the final manuscript.

## Data and materials availability

All the data, code, results produced in the course of this project are available at https://github.com/lijiaxiang111/mollusc-rna-viruses/tree/main. All data and metadata for these sequence records are publicly available and can be accessed via the NCBI (US National Centre for Biotechnology Information) at www.ncbi.nlm.nih.gov. For specific sequence IDs and result files, please refer to the supplementary tables.

## Supplementary Materials

### Supplementary Figures

**Fig. S1.**
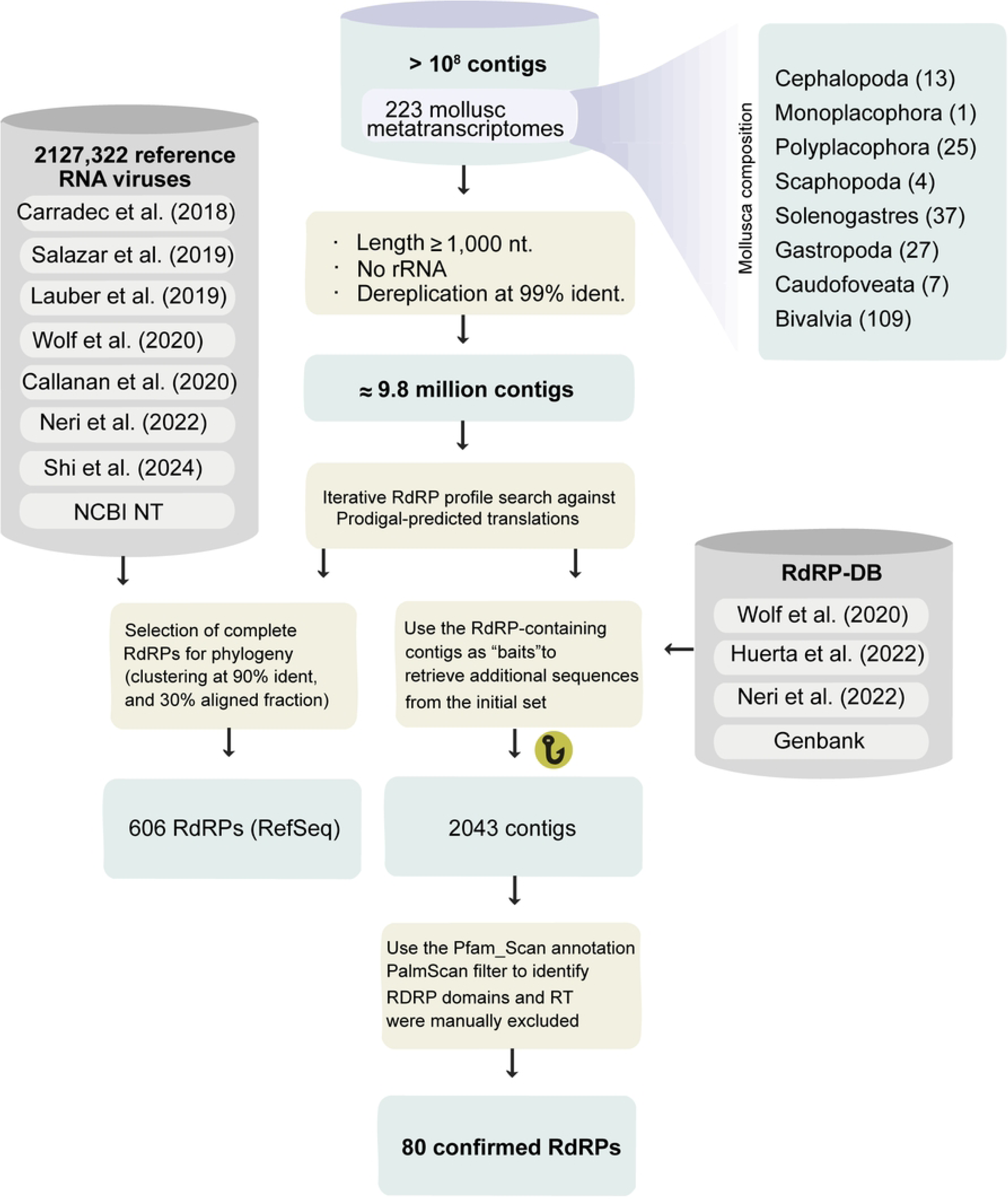
RNA virus discovery pipeline.

**Fig. S2.**
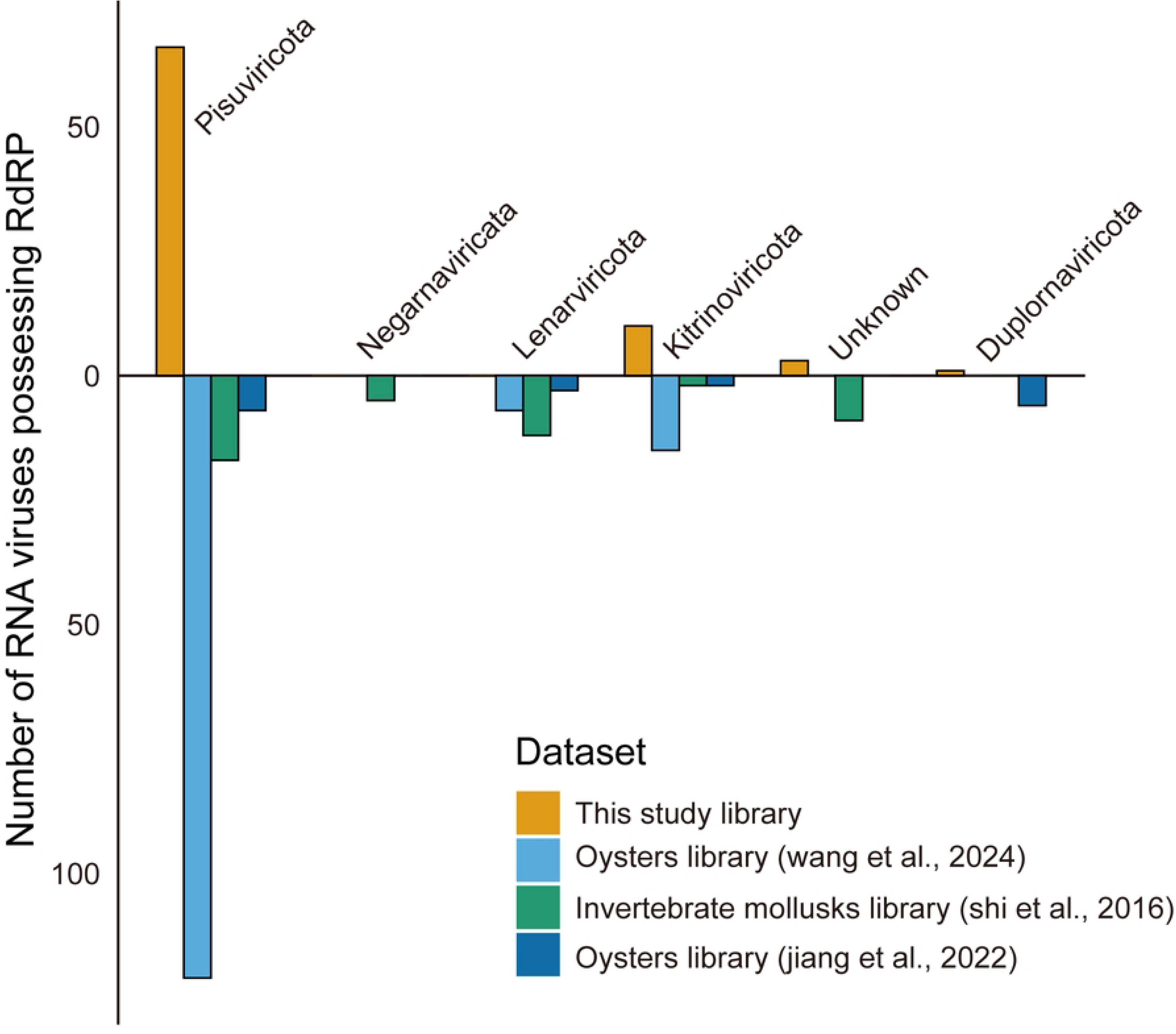
Comparison of RNA virus databases in Molluscs. Standardization of the number of viruses in each library. The main classifications are displayed above the bar chart.

**Fig. S3.**
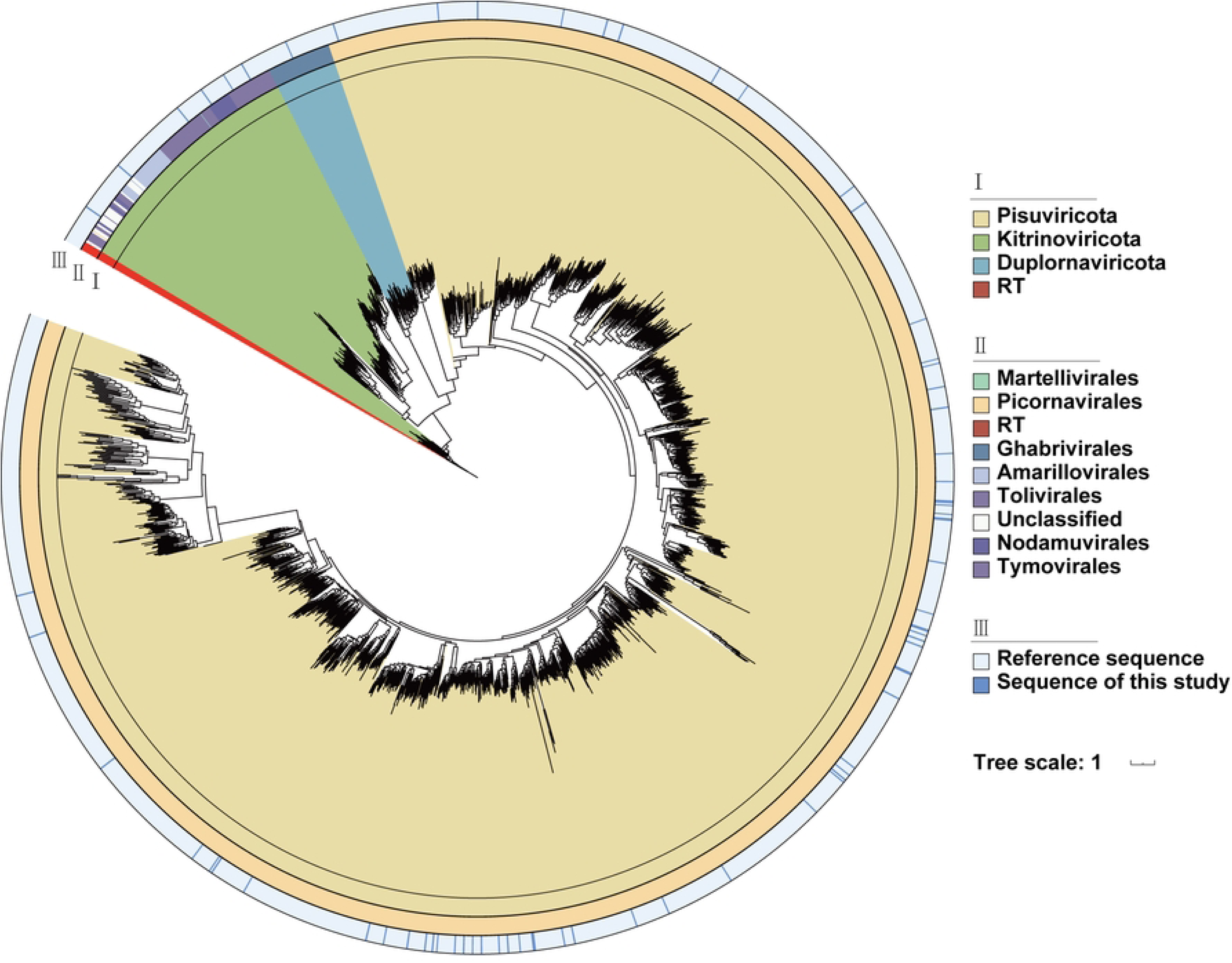
Phylogenetic analysis of RdRP based on phylum and order levels revealed taxonomic diversity of RNA viral contigs in mullusc samples. The phylogenetic tree was visualized using iTOL (https://itol.embl.de/) using reverse transcriptase as an outgroup and as the root of the maximum likelihood tree. The inner ring (Ⅰ) indicates the annotated phyla, the middle ring (Ⅱ) indicates the annotated order, the outermost ring (Ⅲ) indicates the source of the sequence, and the light blue indicates that the virus sequence is from this study.

**Fig. S4.**
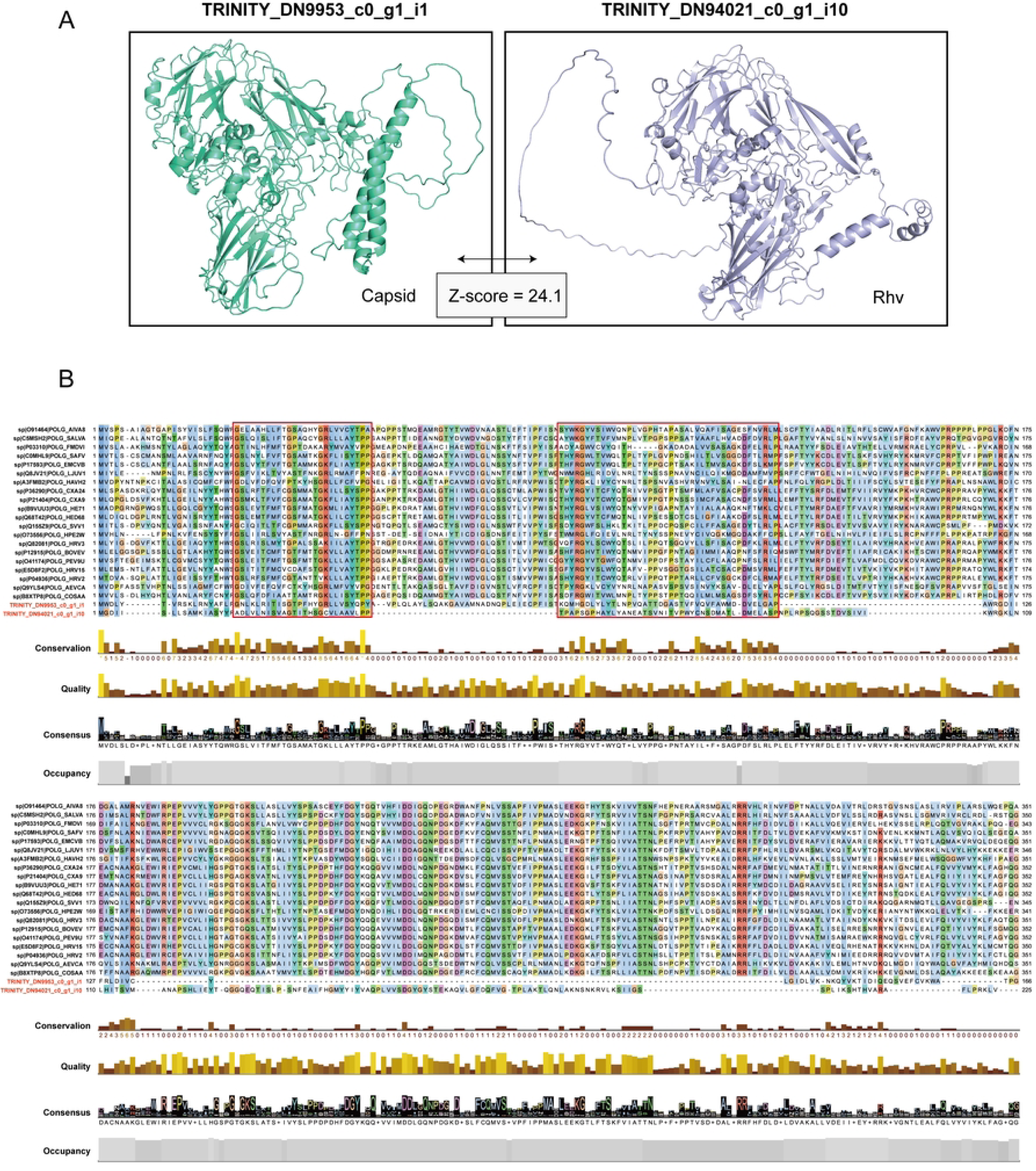
The structural schematic diagram of the capsid carried by *Picornaviridae*. (A) Three-dimensional structure prediction of capsid proteins. The Z-score indicates similarity. (B) The primary structure diagram of the capsid protein of the reviewed Picronaviridae family, arranged based on amino acid conservation. The red box highlights the designated conserved region.

**Fig. S5.**
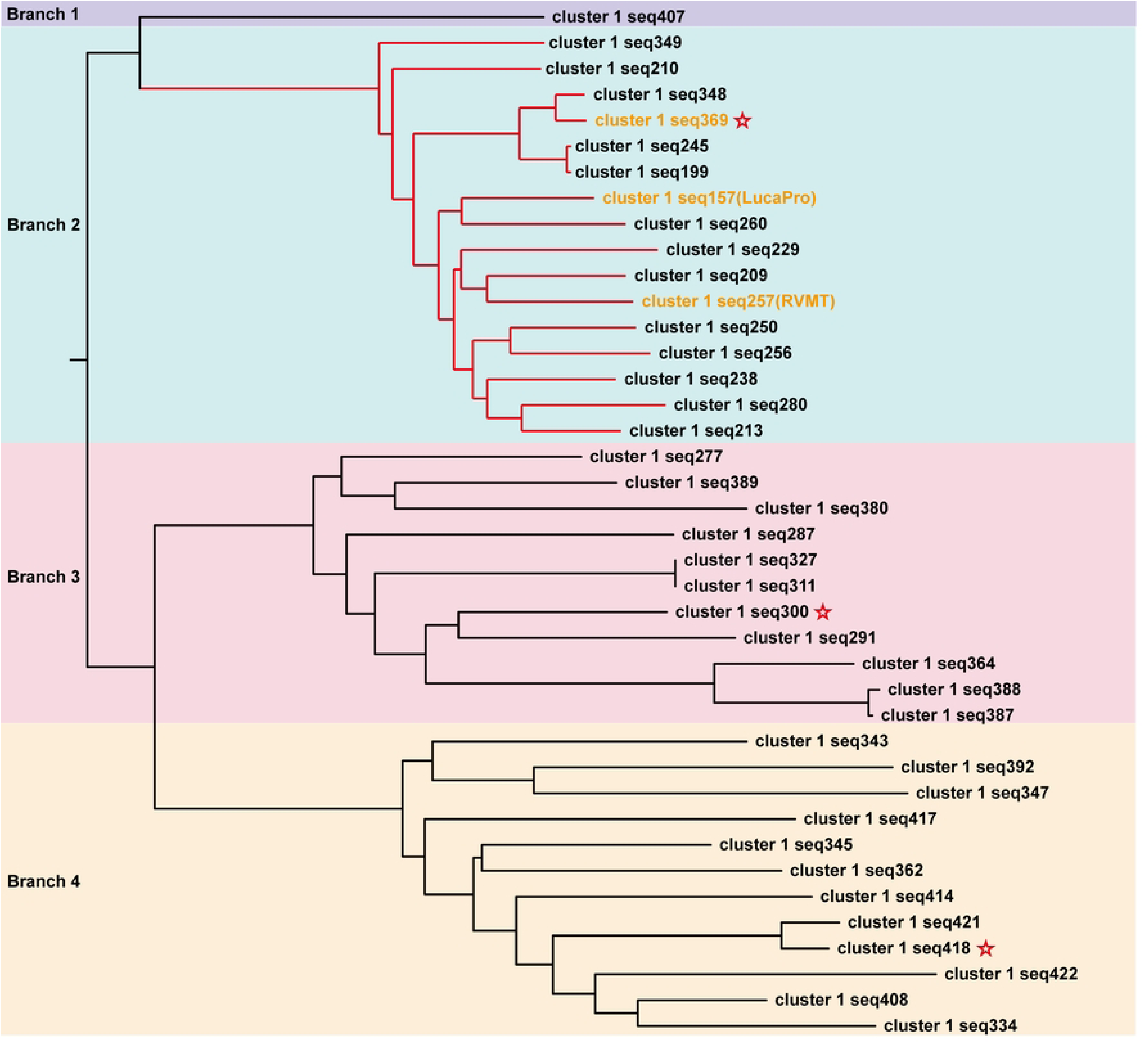
The evolutionary tree branch of *Pisuviricota* was utilized for the analysis of protein characteristics in RNA viral contigs. Red asterisk: Indicates the representative RdRP sequence of this study; Orange label: Denotes the RdRP sequence used for protein characteristic analysis; Different background colors represent distinct branch modules.

**Fig. S6.**
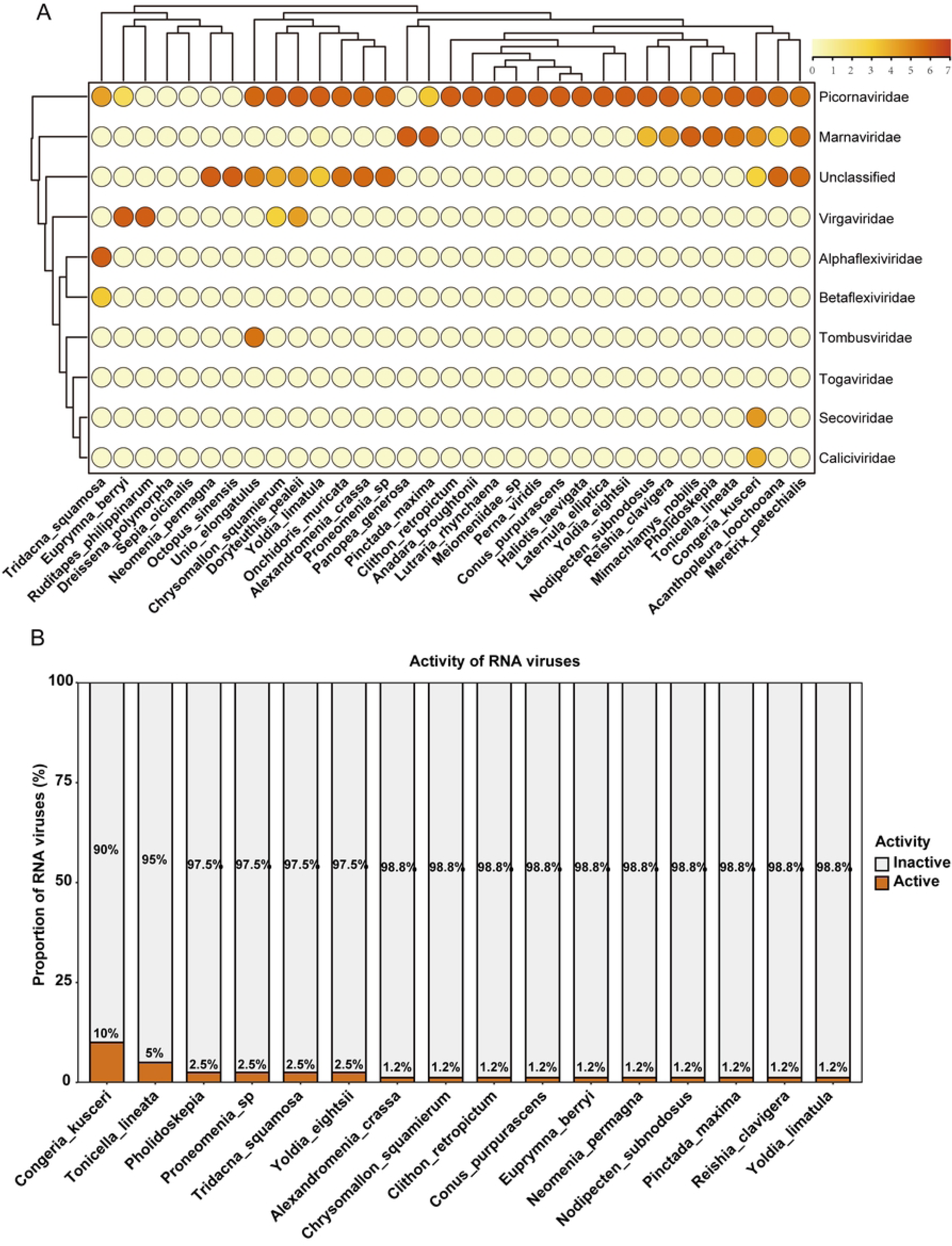
The relative abundance and transcriptional activity of mollusc-associated RNA viruses. (A) The relative abundance of viruses identified in the mollusc determined based on the log10 transformed TPM. (B) Transcriptional activity is measured by the absolute expression level of genes in metaT (TPM/RPKM) combined with the coverage threshold.

## References

1. El-Bawab, F., Chapter 11 - Phylum Mollusca, in Invertebrate Embryology and Reproduction, F. El-Bawab, Editor. 2020, Academic Press. p. 713-813.

2. Kocot, K.M., et al., Phylogenomics reveals deep molluscan relationships. Nature, 2011. 477(7365): p. 452-6.

3. Sun, J., et al., Adaptation to deep-sea chemosynthetic environments as revealed by mussel genomes. Nat Ecol Evol, 2017. 1(5): p. 121.

4. Albertin, C.B., et al., The octopus genome and the evolution of cephalopod neural and morphological novelties. Nature, 2015. 524(7564): p. 220-4.

5. Li, Y., et al., Adaptive Bird-like Genome Miniaturization During the Evolution of Scallop Swimming Lifestyle. Genomics Proteomics Bioinformatics, 2022. 20(6): p. 1066–1077.

6. Young, N.D., et al., Nuclear genome of Bulinus truncatus, an intermediate host of the carcinogenic human blood fluke Schistosoma haematobium. Nat Commun, 2022. 13(1): p. 977.

7. Mushegian, A.R., Are There 1031 Virus Particles on Earth, or More, or Fewer? Journal of Bacteriology, 2020. 202(9): p. 10.1128/jb.00052-20.

8. Call, L., S. Nayfach, and N.C. Kyrpides, Illuminating the Virosphere Through Global Metagenomics. Annu Rev Biomed Data Sci, 2021. 4: p. 369–391.

9. Camargo, A.P., et al., IMG/VR v4: an expanded database of uncultivated virus genomes within a framework of extensive functional, taxonomic, and ecological metadata. Nucleic Acids Research, 2022. 51(D1): p. D733–D743.

10. Shi, M., et al., Redefining the invertebrate RNA virosphere. Nature, 2016. 540(7634): p. 539-543.

11. Oyanedel, D., et al., Cooperation and cheating orchestrate Vibrio assemblages and polymicrobial synergy in oysters infected with OsHV-1 virus. Proceedings of the National Academy of Sciences, 2023. 120(40): p. e2305195120.

12. Renault, T. and B. Novoa, Viruses infecting bivalve molluscs. Aquatic Living Resources, 2004. 17(4): p. 397–409.

13. Elston, R.A. and M.T. Wilkinson, Pathology, management and diagnosis of oyster velar virus disease (OVVD). Aquaculture, 1985. 48(3): p. 189–210.

14. Zhu, P., et al., Novel RNA viruses in oysters revealed by virome. Imeta, 2022. 1(4): p. e65.

15. Wu, S., et al., A comprehensive RNA virome identified in the oyster Magallana gigas reveals the intricate network of virus sharing between seawater and mollusks. Microbiome, 2024. 12(1): p. 263.

16. Wolf, Y.I., et al., Doubling of the known set of RNA viruses by metagenomic analysis of an aquatic virome. Nature Microbiology, 2020. 5(10): p. 1262–1270.

17. Liu, F., et al., MolluscDB 2.0: a comprehensive functional and evolutionary genomics database for over 1400 molluscan species. Nucleic Acids Res, 2025. 53(D1): p. D1075–d1086.

18. Steinegger, M. and J. Söding, MMseqs2 enables sensitive protein sequence searching for the analysis of massive data sets. Nature Biotechnology, 2017. 35(11): p. 1026–1028.

19. Hyatt, D., et al., Prodigal: prokaryotic gene recognition and translation initiation site identification. BMC Bioinformatics, 2010. 11(1): p. 119.

20. Mistry, J., et al., Challenges in homology search: HMMER3 and convergent evolution of coiled-coil regions. Nucleic Acids Research, 2013. 41(12): p. e121–e121.

21. Benson, D.A., et al., GenBank. Nucleic Acids Res, 2013. 41(Database issue): p. D36–42.

22. Neri, U., et al., Expansion of the global RNA virome reveals diverse clades of bacteriophages. Cell, 2022. 185(21): p. 4023–4037.e18.

23. Zayed, A.A., et al., Cryptic and abundant marine viruses at the evolutionary origins of Earth’s RNA virome. Science, 2022. 376(6589): p. 156-162.

24. Wolf, Y.I., et al., Doubling of the known set of RNA viruses by metagenomic analysis of an aquatic virome. Nat Microbiol, 2020. 5(10): p. 1262–1270.

25. Rozewicki, J., et al., MAFFT-DASH: integrated protein sequence and structural alignment. Nucleic Acids Res, 2019. 47(W1): p. W5–w10.

26. Mistry, J., et al., Pfam: The protein families database in 2021. Nucleic Acids Res, 2021. 49(D1): p. D412–d419.

27. Babaian, A. and R. Edgar, Ribovirus classification by a polymerase barcode sequence. PeerJ, 2022. 10: p. e14055.

28. Callanan, J., et al., Expansion of known ssRNA phage genomes: From tens to over a thousand. Science Advances, 2020. 6(6): p. eaay5981.

29. Lauber, C., et al., Discovery of highly divergent lineages of plant-associated astro-like viruses sheds light on the emergence of potyviruses. Virus Res, 2019. 260: p. 38–48.

30. Hou, X., et al., Using artificial intelligence to document the hidden RNA virosphere. Cell, 2024. 187(24): p. 6929–6942.e16.

31. Salazar, G., et al., Gene Expression Changes and Community Turnover Differentially Shape the Global Ocean Metatranscriptome. Cell, 2019. 179(5): p. 1068–1083.e21.

32. Zheng, K., et al., VITAP: a high precision tool for DNA and RNA viral classification based on meta-omic data. Nature Communications, 2025. 16(1): p. 2226.

33. Camargo, A.P., et al., Identification of mobile genetic elements with geNomad. Nat Biotechnol, 2024. 42(8): p. 1303–1312.

34. Buchfink, B., C. Xie, and D.H. Huson, Fast and sensitive protein alignment using DIAMOND. Nature Methods, 2015. 12(1): p. 59–60.

35. Enright, A.J., S. Van Dongen, and C.A. Ouzounis, An efficient algorithm for large-scale detection of protein families. Nucleic Acids Research, 2002. 30(7): p. 1575–1584.

36. Steinegger, M., et al., HH-suite3 for fast remote homology detection and deep protein annotation. BMC Bioinformatics, 2019. 20(1): p. 473.

37. Capella-Gutiérrez, S., J.M. Silla-Martínez, and T. Gabaldón, trimAl: a tool for automated alignment trimming in large-scale phylogenetic analyses. Bioinformatics, 2009. 25(15): p. 1972–3.

38. Nguyen, L.-T., et al., IQ-TREE: A Fast and Effective Stochastic Algorithm for Estimating Maximum-Likelihood Phylogenies. Molecular Biology and Evolution, 2014. 32(1): p. 268–274.

39. Lin, Z., et al., Evolutionary-scale prediction of atomic-level protein structure with a language model. Science, 2023. 379(6637): p. 1123-1130.

40. van Kempen, M., et al., Fast and accurate protein structure search with Foldseek. Nat Biotechnol, 2024. 42(2): p. 243–246.

41. Bastian, M., S. Heymann, and M. Jacomy. Gephi: an open source software for exploring and manipulating networks. in Proceedings of the international AAAI conference on web and social media. 2009.

42. Chen, G., J. Jiang, and Y. Sun, RNAVirHost: a machine learning-based method for predicting hosts of RNA viruses through viral genomes. Gigascience, 2024. 13.

43. Langmead, B. and S.L. Salzberg, Fast gapped-read alignment with Bowtie 2. Nat Methods, 2012. 9(4): p. 357–9.

44. Li, H., et al., The Sequence Alignment/Map format and SAMtools. Bioinformatics, 2009. 25(16): p. 2078–9.

45. Coclet, C., et al., Virus diversity and activity is driven by snowmelt and host dynamics in a high-altitude watershed soil ecosystem. Microbiome, 2023. 11(1): p. 237.

46. Letunic, I. and P. Bork, Interactive Tree of Life (iTOL) v6: recent updates to the phylogenetic tree display and annotation tool. Nucleic Acids Res, 2024. 52(W1): p. W78–w82.

47. Wu, J., et al., Structural basis of transition from initiation to elongation in de novo viral RNA-dependent RNA polymerases. Proc Natl Acad Sci U S A, 2023. 120(1): p. e2211425120.

48. Du, X., The cellular RNA-dependent RNA polymerases in plants. New Phytol, 2024. 244(6): p. 2150–2155.

49. Rath, D., et al., The CRISPR-Cas immune system: biology, mechanisms and applications. Biochimie, 2015. 117: p. 119–28.

50. Koonin, E.V., et al., Global Organization and Proposed Megataxonomy of the Virus World. Microbiology and Molecular Biology Reviews, 2020. 84(2): p. 10.1128/mmbr.00061-19.

51. Culley, A.I., A.S. Lang, and C.A. Suttle, Metagenomic analysis of coastal RNA virus communities. Science, 2006. 312(5781): p. 1795-8.

52. Carella, F., et al., A widespread picornavirus affects the hemocytes of the noble pen shell (Pinna nobilis), leading to its immunosuppression. Front Vet Sci, 2023. 10: p. 1273521.

53. Zhuo, R., et al., Molecular Epidemiology of Human Sapovirus among Children with Acute Gastroenteritis in Western Canada. J Clin Microbiol, 2021. 59(10): p. e0098621.

54. Glass, R.I., U.D. Parashar, and M.K. Estes, Norovirus gastroenteritis. N Engl J Med, 2009. 361(18): p. 1776–85.

55. Dos Santos, N.L., et al., Occurrence and Molecular Characterization of Human Astrovirus and Hepatitis A Virus in Bivalve Mollusks Marketed in Tourist Cities in Rio de Janeiro, Brazil. Food Environ Virol, 2025. 17(2): p. 23.

56. Wu, R., et al., RNA Viruses Linked to Eukaryotic Hosts in Thawed Permafrost. mSystems, 2022. 7(6): p. e0058222.

